# Development of reduced gluten wheat enabled by determination of the genetic basis of the *lys3a* low hordein barley mutant

**DOI:** 10.1101/354548

**Authors:** Charles P. Moehs, William J. Austill, Aaron Holm, Tao A. G. Large, Dayna Loeffler, Jessica Mullenberg, Patrick S. Schnable, Wayne Skinner, Jos van Boxtel, Liying Wu, Cate McGuire

**Author notes:** present address: Phytelligence 221 1^st^ Ave W. Suite 350, Seattle, WA 98119. present address: Mailbox 132, Mudd Building, Rm. 121, MC5080, 333 Campus Dr. Stanford, CA 94305. corresponding authors: Charles P. Moehs,; Jos van Boxtel.

## Abstract

Celiac disease is the most common food-induced enteropathy in humans with a prevalence of approximately 1% world-wide [1]. It is induced by digestion-resistant, proline- and glutamine-rich seed storage proteins, collectively referred to as “gluten,” found in wheat. Related prolamins are present in barley and rye. Both celiac disease and a related condition called non-celiac gluten sensitivity (NCGS) are increasing in incidence [2] [3]. This has prompted efforts to identify methods of lowering gluten in wheat, one of the most important cereal crops. Here we used BSR-seq (Bulked Segregant RNA-seq) and map-based cloning to identify the genetic lesion underlying a recessive, low prolamin mutation (*lys3a*) in diploid barley. We confirmed the mutant identity by complementing the *lys3a* mutant with a transgenic copy of the wild type barley gene and then used TILLING (Targeting Induced Local Lesions in Genomes) [4] to identify induced SNPs (Single Nucleotide Polymorphisms) in the three homoeologs of the corresponding wheat gene. Combining inactivating mutations in the three sub-genomes of hexaploid bread wheat in a single wheat line lowered gliadin and low molecular weight glutenin accumulation by 50-60% and increased free and protein-bound lysine by 33%. This is the first report of the combination of mutations in homoeologs of a single gene that reduces gluten in wheat.

---

Gluten is the common name for a complex mixture of approximately 100 proline- and glutamine-rich seed storage proteins found in wheat endosperm. In hexaploid bread wheat, in particular, gluten is responsible for the unique viscoelastic properties of dough used for making leavened bread. Related prolamins known as hordeins in barley and secalins in rye, but lacking the viscoelastic properties of wheat gluten, are found in the endosperms of these related cereals. Among gluten proteins, the most important for the elasticity and strength of bread dough are the polymeric high molecular weight glutenins [5]; monomeric gluten proteins, primarily gliadins, which are classified based on size, charge and repetitive amino acid sequences into α, β, γ and ω types, contribute to the viscosity of gluten [6] [7].

Gluten is the causative factor of celiac disease, the most common food-induced autoimmune enteropathy in humans [1]. Celiac disease is found primarily in individuals expressing human leukocyte antigen (HLA) alleles DQ2 and/or DQ8. Immunodominant peptide epitopes have been found in α gliadins [8] and in ω gliadins of wheat and in C hordeins of barley [9]. Additional epitopes are also found in LMW and HMW-glutenins, gamma-gliadins from wheat and B and gamma hordeins from barley [10] [9]. Celiac disease patients exhibit a range of intestinal and extra-intestinal symptoms of which the most common is flattening of the villi of the small intestine, which can lead to poor absorption of nutrients and other pathological consequences. The only current treatment is a life-long avoidance of all gluten in the diet, which can be challenging given the prevalent use of gluten in diverse processed food products.

Given gluten’s negative consequences for celiac patients, efforts have been made to lower gluten by combining gliadin deletion mutants [11] or by using RNAi transgenic methods in wheat [12] [13] [14], as well as by conventional breeding of known low hordein mutants in barley [15]. One such mutant is *lys3a*, a low hordein barley that is also known as Risø 1508 or sex3c (shrunken endosperm xenia) [16]. It was induced by ethylenimine mutagenesis of the two-rowed spring malting variety, Bomi [17] close to 50 years ago in a program of barley mutagenesis and screening for increased lysine at the Risø agricultural experiment station in Roskilde, Denmark [18, 19]. The *lys3a* mutant is almost completely lacking in class C hordeins, and accumulates considerably reduced amounts of several B class hordeins, while having a 45% increase in the accumulation of free and protein bound lysine in the seed compared to the parental control [20] [21] [22]. These phenotypes are the result of a single recessive allele [23] that has been variously mapped to barley chromosome 5H [24] [25] [16], or, more recently, to chromosome 1H [26] [27], but the underlying mutant gene is unknown.

We undertook to identify the *lys3a* mutant gene in barley using bulked segregant RNA-seq (BSR-seq) [28] and genetic fine mapping with the objective to determine if an analogous low gluten wheat could be developed by inactivation of the homoeologs in wheat. In the following, we show that the *lys3a* mutation is due to a missense allele in a Domain of One Finger (DOF) zinc finger transcription factor known as barley prolamin-box binding factor (BPBF) [29]. Transformation of the *lys3a* barley mutant with the wild type BPBF gene restored hordein levels, which confirms the fact that a mutation in BPBF is responsible for the *lys3a* phenotype. Finally, we show that mutating the wheat homoeologs of BPBF, the wheat prolamin-box binding factors (WPBFs), using the non-transgenic TILLING method, results in a reduced gluten, high lysine wheat. This is the first demonstration of a reduced gluten, high lysine wheat developed without the use of transgenic methods.

## Methods

### Genetic Crosses and Hordein Extraction

The original *lys3a* mutant and its parent, Bomi, were obtained from the USDA’s National Small Grains Germplasm Collection in Aberdeen, Idaho (https://www.ars.usda.gov/pacific-west-area/aberdeen-id/small-grains-and-potato-germplasm-research/docs/national-small-grains-collection/). *Lys3a* was crossed to cultivar Bowman as well as to Lysiba, a cultivar developed with the original *lys3a* allele. To evaluate the presence of the recessive *lys3a* allele in seeds, F2 and subsequent generation seeds were cut in half and hordeins from endosperms were extracted according to [22] [30] and evaluated by SDS-PAGE. The embryo halves of the seeds were saved for planting.

### Plant Material

Hexaploid wheat (cv. Express) with mutations in the WPBF (wheat prolamin-box binding factor) gene in each of the A, B and D sub-genomes were crossed to each other and to the parental cultivar. Different genetic classes were identified by KASP genotyping (https://www.lgcgroup.com/products/kasp-genotyping-chemistry/#.WcQPHsZrzZ4) using probes developed to specific SNPs and genomic DNA isolated from seedling leaf tissue. To study restoration of hordein accumulation in the *lys3a* mutant through complementation by BPBF expression, *lys3a*-containing cultivar Lysiba was used for transformation.

### Protein Extraction

Prolamin proteins were extracted from mature wheat and barley seeds as described [31]. Wheat seeds (1 g) were ground to a fine powder using a Retsch Model 301 mixer mill. Due to segregation of alleles in T1 barley transgenic for the BPBF gene, twelve individual seeds were separately ground, extracted and analyzed.

For protein extraction, 40 mg of ground seed was extracted with 1 mL of 60 % isopropanol and 2% beta-mercaptoethanol in 50 mM Tris-Borate buffer, pH 8.0, by heating at 60 °C for 1 hour with mixing. Extracted samples were centrifuged (18,000 *g* × 10 min) and a 0.525 mL aliquot added to 1.5 mL of 20% NaCl and stored in a −20°C freezer overnight to precipitate prolamin proteins. Precipitated proteins were pelleted at 18,000 *g* for 2 min, the supernatant decanted, and residual solution removed by vacuum evaporation using a SpeedVac concentrator. Protein pellets were then dissolved in Sigma Total Plant Protein Reagent Type 4 containing 1% Sigma Plant Protease Inhibitor Cocktail. The protein concentrations were determined using either the Pierce 660 nm protein assay or the Qubit protein assay (Thermo Fisher Scientific).

### Protein Gel Electrophoresis

The extracted prolamins were denatured and reduced with Novex Tris-Glycine SDS Sample Buffer and Reducing Agent (Thermo Fisher Scientific) and heating to 70 °C for 10 min. SDS-PAGE was performed using Invitrogen WedgeWell^™^ Tris-Glycine gels and Bio-Rad Kaleidoscope Precision Plus MW Standards with Novex Tris-Glycine running buffer. Developed gels were fixed in 50% methanol/10% acetic acid solution for 15 min and stained for 3 hr using Colloidal Blue Staining Kit (Thermo Fisher Scientific). Gel images were obtained using a BioRad ChemiDoc XRS+ system with Image Lab software.

### Liquid Chromatography Mass Spectrometry

Gluten proteins in SDS-PAGE gel bands were identified by data-dependent LC-MS^n^ [32]. Excised gel bands were diced, destained, reduced and alkylated using the Pierce in-gel protein digestion kit prior to mass spectrometric analysis [33]. In-gel digestion of proteins with chymotrypsin (Promega) for 16 hr at 37° C followed by Trypsin (Promega) for 6 hr. Peptides were separated on an ACE C18 column (0.3 × 150 mm, Advanced Chromatography Technologies) in a 5 to 55% gradient of acetonitrile in 0.1% formic acid. Spectra were obtained on a LCQ Deca XP plus mass spectrometer (Thermo Scientific) using one survey MS scan (350 to 2000 Da) followed by four data-dependent MS/MS scans of the most abundant ions in the MS survey scan. Parameters for data-dependent MS/MS included a default charge state of 2, an isolation width of 2.0 Da and collision energy of 35% for collision induced decay (CID) of ions having abundance greater than 1 × 10^5^.

Mass spectral data were processed using the GPM manager application (GPM extreme edition, v. 2.2.1.0, Beavis Informatics Ltd, Manitoba, Canada) and Scaffold^™^ software (v. 3.00.03, Proteome Software Inc., Portland, OR) with comparison to proteins in the UniProtKB *Triticum aestivum* database (downloaded on May 25, 2016) to which the sequence of common contaminants were appended (e.g. human keratins, trypsin, BSA, and others in the cRAP database from GPM) prior to reverse concatenation using a Perl script (provided by Dr. Brett Phinney, UC Davis Genome Center Proteomics Core Facility). To assert whether a protein is present in a given sample, a minimum of two peptides with greater than 80% probability of being correctly identified was required as well as a minimum 99% probability of protein presence. Probabilities were assigned by the Peptide Prophet and Protein Prophet algorithms within the Scaffold bioinformatics software [34] [35] [36] [37].

### Amino Acid Analysis

Total amino acid content of 1 g. powdered whole wheat seed was determined by Covance Labs (Madison, WI) using automated pre-column derivatization of hydrolysates followed by HPLC analysis according to the methods of [38] and [39] and included the analysis of cysteine [40]. The analyses were conducted on three biological replicates.

### BSR-seq

BSR-seq was conducted according to [28]. RNA was isolated from three independent biological replicate pools of wild type and three independent biological replicate pools of homozygous mutant tissue. Each pool consisted of 7-day old etiolated shoot and root tissue from between 20-30 germinated F3 seeds, each derived from a separate F2 seed from a *lys3a* × Bowman cross. In the mutant pool, F2 recessive mutant seeds were identified by half-seed hordein analysis as described above. To ensure that individuals in the wild type pool were not segregating for the recessive *lys3a* allele, F2 embryos whose endosperm half-seed hordein analysis indicated a wild type phenotype were planted and half-seed hordein analysis was conducted on approximately 20 F3 seeds from each wild type F2. In the wild type pool, only tissue from germinated F3 seeds was used, none of whose siblings had any mutant segregants.

### Marker Development, Fine Mapping and Identification of BPBF

Using the sequence context information from SNPs that were exclusively linked to the mutant pool, KASP probes (LGC, https://www.lgcgroup.com/) were designed and used to genotype individuals from a larger mapping population. Hordein extracts from endosperm half-seeds of 1814 seeds were evaluated to identify recessive *lys3a* mutants. DNA was isolated from seedling leaf tissue grown from the germinated embryos. Following fine-mapping, PCR primers were designed to amplify the BPBF gene. PCR products were amplified from genomic DNA isolated from seedling leaf tissue of Bomi and *lys3a* using a proof-reading polymerase (Phusion polymerase, New England Biolabs) and sequenced.

### Transformation of Barley

Construct pARC1057 (**Figure S1**) was transformed into immature embryos of barley cv. Lysiba, essentially as described [41], with some modifications. The *Agrobacterium* strain used was EHA105 [42], timentin was added to the medium at 200 mg/l to suppress bacterial growth after co-cultivation and transgenic plant tissue was selected by using geneticin at 100 mg/l. After 3 weeks of rooting of shoots in medium containing 25 mg/l kanamycin, T_0_ plantlets were transferred into a mixture of Sunshine #2 and Sunshine #3, in a 1:1 ratio, and grown at day/night periods of 14h at 25°C/10h at 18°C. Expression of the NPTII gene was confirmed by analyzing ground T_0_ leaf material by NPTII immunostrip test (Agdia). Harvested T_1_ seed was planted and grown similarly as described above.

### TILLING of WPBF Homoeologs

TILLING of wheat was conducted according to [43]. Approximately 10,000 individual M2 mutant genomic DNAs from spring wheat cultivar Express were screened. TILLING was conducted both by an acrylamide gel-based screen and by a capillary electrophoresis-based method.

## Results

### Identification of *lys3a* Mutation as BPBF

To gain insight into the molecular mechanisms responsible for the *lys3a* mutant’s effects, we conducted a BSR-seq experiment [28] to identify the underlying lesion. Our BSR-seq experiment was conducted on the F3 progeny of a cross between *lys3a* and Bowman. Separation of the F2 seeds derived from this cross into normal and mutant phenotypes was facilitated by the ease of distinguishing the homozygous recessive mutant from the normal phenotype by analysis of hordeins extracted from endosperm tissue as shown in **Figure 1**. When loading equal fractions of extracted hordeins from normal and mutant endosperms, hordeins from the homozygous *lys3a* mutant parent, including B, C and D hordeins, are not visible on a Coomassie stained SDS-PAGE gel compared to the conventional non-mutant Bowman barley cultivar. Only when the gel is overloaded 4-fold as shown in panel B do hordeins become visible in the mutant. This facile SDS PAGE screen of F2 half-seeds was used to identify wild type and homozygous mutant progeny of the cross. The tissue bulks used for RNA isolation were derived from 25-35 individual F3 seedlings, each derived from 25-35 independent F2s determined to be either homozygous wild type or homozygous mutant. To ensure that the wild type F3 bulks were derived from homozygous wild type F2 seeds, we ran SDS PAGE gels of hordeins extracted from F3 progeny of the wild type F2s and discarded those that segregated mutants. Illumina sequencing of cDNA derived from RNA isolated from three biological replicates of both mutant and wild type whole-seedling tissue bulks identified SNPs and gene expression variation tightly linked with the mutation.

**Figure 1.**
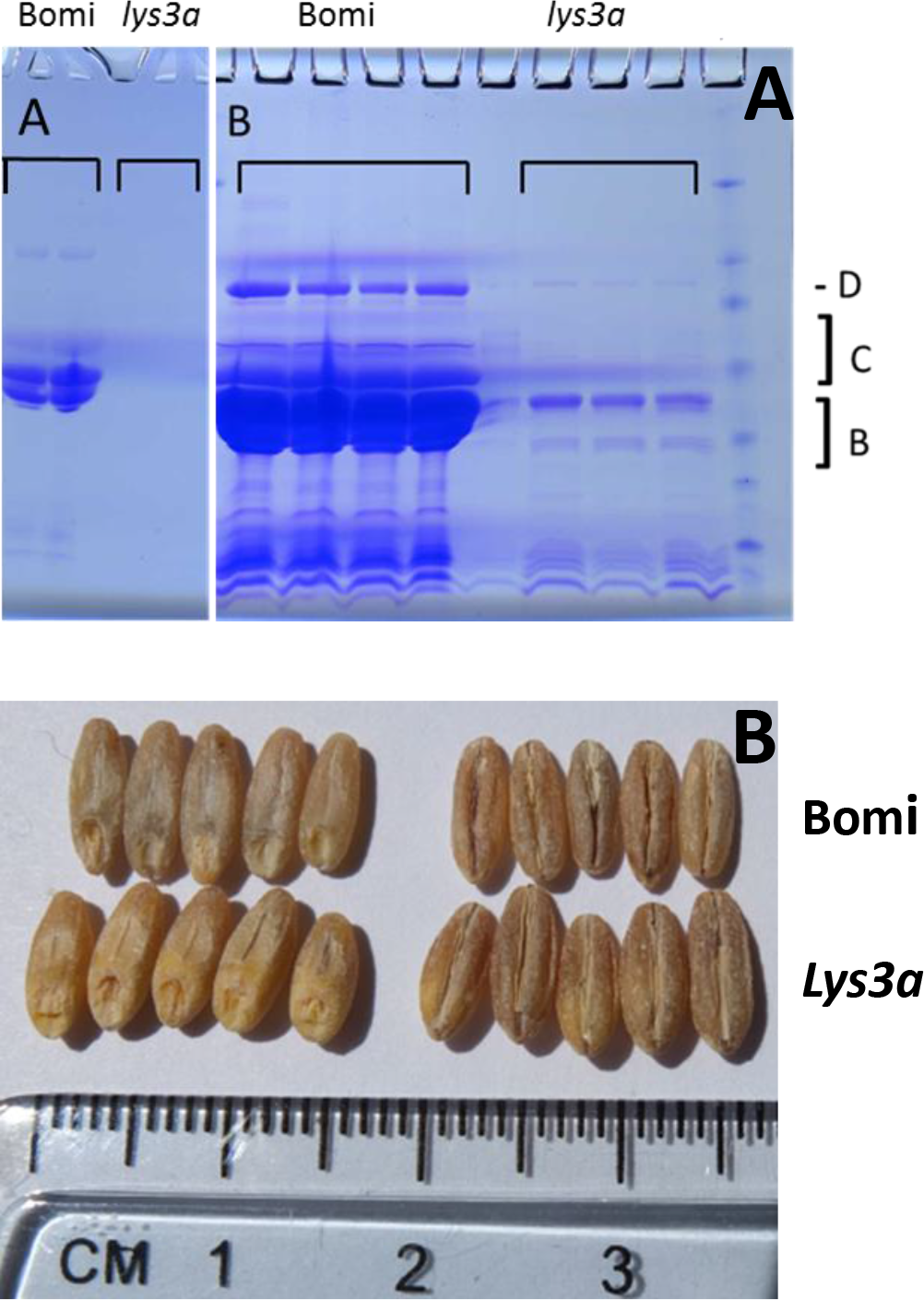
SDS-PAGE of Bomi and *lys3a* barley endosperm hordeins (A) and appearance of seeds (B) (A) 4-20% gradient SDS-PAGE gel electrophoresis experiment in which endosperm hordeins of parental cultivar Bomi, and the *lys3a* mutant derived from it, are shown. Equivalent fractions of extracted hordeins from both are shown. Panel B represents a 4 four-fold overloading of the hordeins compared to panel A. D refers to the position of the D hordein, C shows the location of the C hordeins and B indicates the location of the B hordeins on the gel. (B) Image of seeds of Bomi and *lys3a* mutant derived from Bomi (husks removed) (top and bottom rows respectively).

This experiment allowed us to definitively link the *lys3a* mutant to an approximately 7 cM interval in the pericentromeric region of chromosome 5H (**Figure 2**). Utilizing the first (preliminary) version of the barley genome assembly [44], this area appears to encompass a physical distance of approximately 200 Mbp of DNA in a region of suppressed recombination. We generated 25 KASP genotyping probes (**Table S1**) (https://www.lgcgroup.com/) from some of the 123 SNPs linked to the mutant on chromosome 5H and mapped them in a larger mapping population of 1814 individuals. Half-seed hordein analysis of all seeds identified homozygous mutant individuals, while KASP genotyping identified co-segregating markers. Two markers, approximately 1 Mbp apart, were in absolute linkage with the mutant phenotype with no recombinants (**Figure S2**). Examination of this region of barley chromosome 5H using Ensembl Barley (http://plants.ensembl.org/Hordeum_vulgare/Info/Index) [45] and BarleyMap (http://floresta.eead.csic.es/barleymap/) [46] [47] identified approximately 30 HC (high confidence) and 30 LC (low confidence) genes. We examined this region for putative regulatory genes such as transcription factors and determined that a LC gene was identical to the known BPBF gene. Sequencing of this gene in the *lys3a* and Bomi backgrounds identified a novel SNP in *lys3a* compared to the Bomi parental line. This SNP resulted in a missense mutation in the DOF-DNA binding zinc finger, and we considered it to be a likely candidate for the barley *lys3a* mutant. In *lys3a*, an A to T transversion at nucleotide 173 (numbering beginning with the <u>A</u>TG start codon) leads to a missense mutation in which glutamine at amino acid 58, is replaced by leucine (**Figure 3**). Barley BPBF was cloned in 1998 [29] on the basis of the homology of its DOF DNA binding domain to the previously isolated *Zea mays* DOF transcription factor that was named Prolamin-box Binding Factor since it binds to a highly conserved seven nucleotide motif called the P-box (Prolamin box) that is found in the promoters of multiple seed storage protein genes [48]. Mena et al., (1998) also cloned a cDNA of a wheat homolog of the barley DOF transcription factor. Subsequent research showed that the wheat homolog indeed plays a role in the transcription of gliadin genes [49], and other researchers cloned and sequenced the three genomic homoeologs of hexaploid bread wheat WPBF [50].

**Figure 2.**
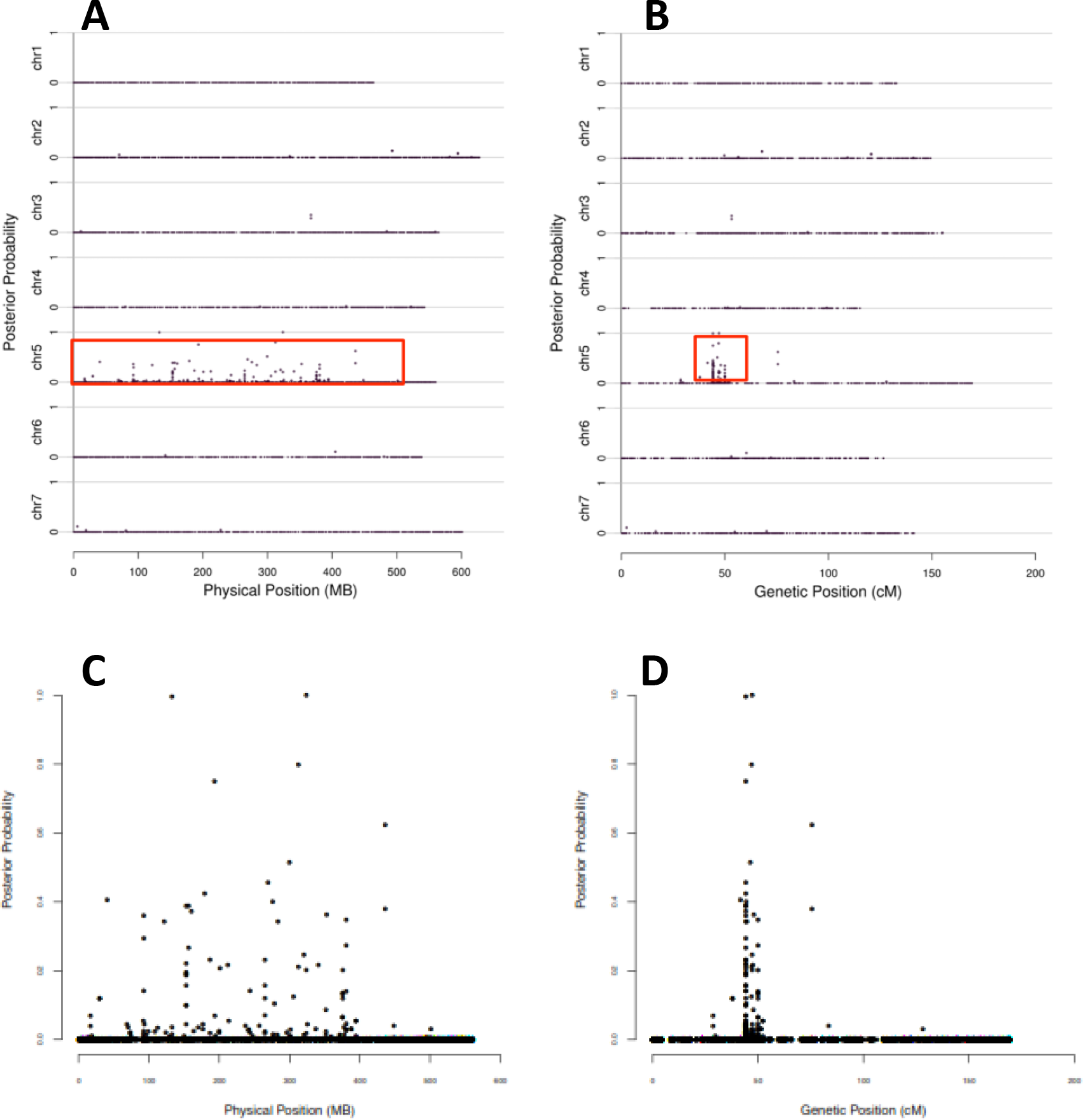
BSR-seq localizes*lys3a* to chromosome 5H. Posterior probabilities of SNPs putatively co-segregating with *lys3a* are depicted on the y axis while physical position (A, C) or genetic position (B, D) are shown on the x axis. A and B show all seven barley chromosomes while C and D depict chromosome 5H only

**Figure 3.**
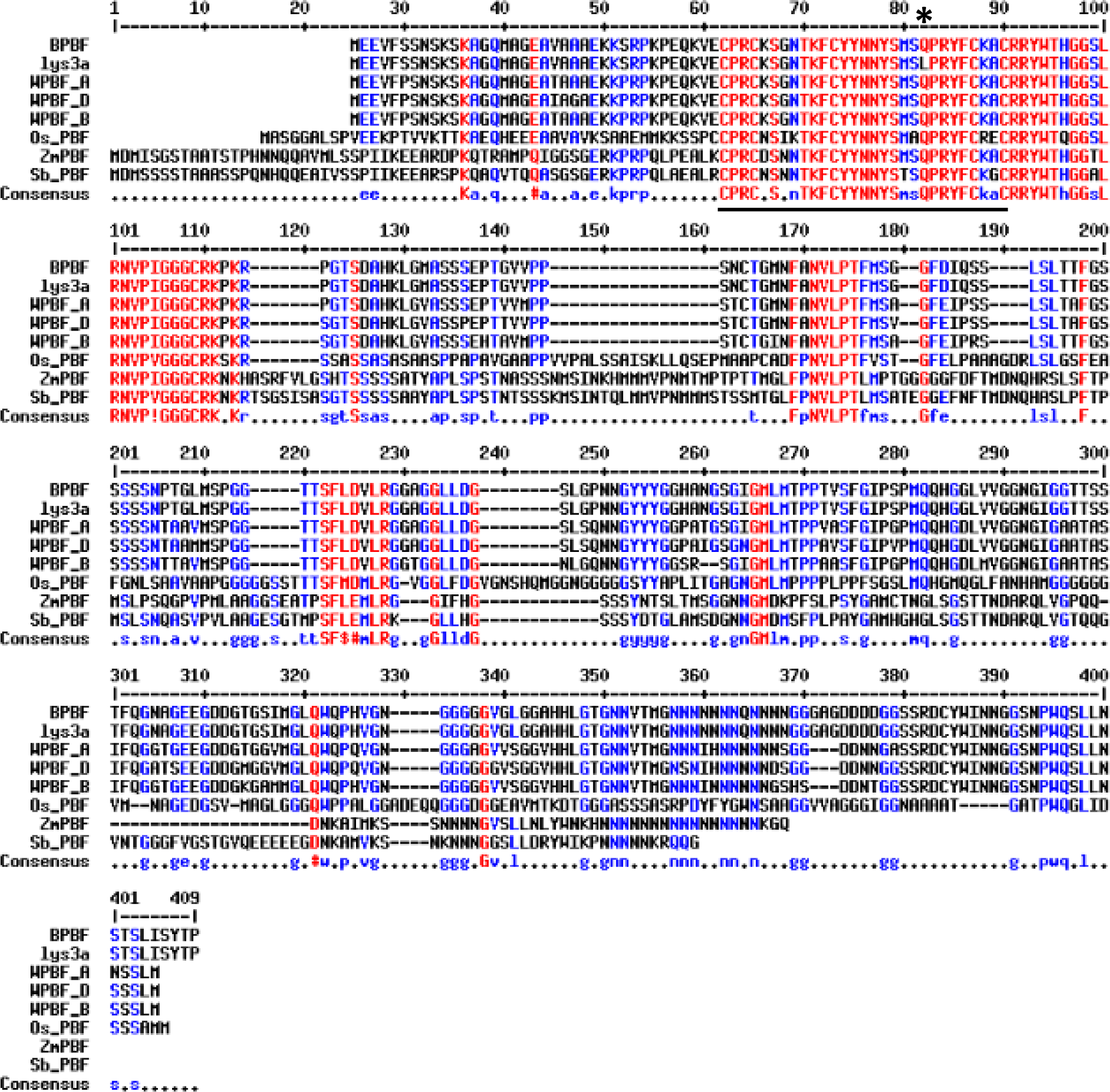
Alignment of cereal PBF proteins. A sequence alignment of the barley, the *lys3a* barley mutant, the wheat A, B, and D sub-genome homoeologs, as well as the rice, maize and sorghum prolamin box binding factor proteins is shown. The zinc finger is underlined and an asterisk denotes the absolutely conserved glutamine, which is mutated to a leucine in the *lys3a* barley mutant. The alignment was made using default parameters of the program multalin [82]. Accession number for rice PBF gene is AK107294 [83], for the maize PBF is AAB70119.1 [48], and for the sorghum PBF is XP_002448852.1.

The glutamine which is altered to leucine in *lys3a* barley is an absolutely conserved residue in the zinc finger DNA binding region of the DOF domain [51] (**Figure 3**), and this alteration from glutamine, an amino acid with a polar side chain, to leucine, an amino acid with a hydrophobic side chain, is predicted to severely negatively affect the DNA binding properties of the DOF domain using the PROVEAN (**<u>Pro</u>**tein **<u>V</u>**ariation **<u>E</u>**ffect **<u>An</u>**alyzer) bioinformatics server [52] This Q>L alteration of the DOF domain in *lys3a* barley was also found upon sequencing the same gene in Lysiba, the barley cultivar developed by outcrossing the original mutant at least three times to unrelated cultivars and selecting at each cross for high lysine segregants. Based on the known properties of BPBF, we hypothesized that BPBF represented a strong candidate for underlying the *lys3a* mutation and we performed additional experiments to verify this hypothesis.

### Transgenic Complementation of the *lys3a* Mutant with the Wild Type BPBF Gene

We transformed the mutant cultivar Lysiba with a construct in which the BPBF gene from the Bomi parent was driven by the strong seed-specific D-hordein gene promoter [53]. This promoter was chosen because it had been previously shown to result in strong endosperm-specific transgene expression and because the D-hordein gene is not itself regulated by the BPBF gene [54]. Of 6 independent transformation events recovered, 2 lines exhibited T1 seeds with increased hordein levels (**Figures S3, S4 and S5**). In these 2 transgenic lines, some seeds exhibited a more than 2-fold overall increase in hordein levels, particularly C-hordein. Transgenic complementation did not restore all missing hordein proteins in these two transgenic lines. This result may be due to differences in the expression pattern of the native BPBF gene compared to the expression pattern of the D-hordein promoter driving expression of the BPBF gene in the transgenic lines.

### Genetic Variation in the Barley BPBF Gene in the USDA Barley Core Germplasm Collection

To assess the degree of genetic variation in the barley PBF gene, and to determine whether any similar alleles that alter the DOF domain in this gene exist in barley germplasm collections, we obtained a set of 186 barley accessions from the USDA’s National Small Grains Germplasm Collection that represent a “mini-core” of the larger 1860 member “iCore” (informative Core) collection of diverse barley germplasm [55]. This 10% subset of the iCore was chosen based on its contribution to the polymorphism information content (PIC) of the larger core set (**Table S2**). Genomic DNA was prepared from seedlings of each of the members of the mini-core and the BPBF gene was PCR amplified using Phusion high-fidelity polymerase and sequenced from all 186 members (list of primers **Table S3**). High quality sequence was obtained from 180 of the 186 members of this mini “iCore”. The sequence extended from 434 bases 5’ of the ATG protein start codon to 580 bases 3’ of the stop codon (2,017 bp total). Twelve distinct haplotypes were identified; seven haplotypes were represented by a single accession and two haplotypes were found in two accessions each (**Table S2** and **Figure S6**). The remaining accessions (169) were represented by three haplotypes: haplotype 1 with a sequence identical to Bomi was the most abundant. It was found in 91 accessions. Haplotype_11, which differed from the sequence of Bowman at a single position, was the second most abundant haplotype. It was present in 63 accessions. The haplotype represented by Bowman (haplotype_9) was found in 15 accessions. The two parents, Bomi and Bowman (haplotype_1 and haplotype_9, respectively), in our mapping population, differ at 10 SNPs in the 999 bp BPBF protein coding sequence, five of which affect the protein sequence (A21V, A154G, S196G, S238N and V266G; **Figure S7**) When analyzed with the PROVEAN program [52], none of these changes is anticipated to alter transcription factor function. Six accessions contained additional SNPs that affect the protein sequence but none of the SNPs (P32L, P91T, G118R, G306D) were found in the zinc finger DNA binding domain (**Figure S7**). Analysis of these variants with the PROVEAN server suggested that the P91T, G118R and G306D alleles were neutral and unlikely to affect BPBF function. The P32L allele, however, was scored as moderately deleterious. Further analysis of the two accessions containing this allele will be required to assess whether it influences hordein accumulation.

A previous study, using a different collection of barley accessions, had previously sequenced two shorter fragments of the BPBF gene in a candidate gene association study of barley endosperm agronomic traits and identified some of the same haplotypes [56]. Consistent with our results, these authors identified SNPs in BPBF associated with crude protein content.

### TILLING of WPBF, the Wheat Homoeologs of the Barley PBF Gene

To determine whether inactivating mutations of the homoeologs of WPBF, the wheat PBF gene, affect wheat seed storage protein accumulation in a manner similar to the *lys3a* allele in barley, we used TILLING in a hexaploid wheat TILLING library containing more than 10,000 individual M2 lines [43] to identify novel alleles in all three wheat sub-genomes. TILLING primers (**Table S3**) were devised that were specific for each of the three genomes of wheat. In total 789 TILLING alleles were found, however due to redundancy in the library, of these 789 alleles, only 488 were unique (numbers of unique mutants shown in parentheses in **Table 1**). In the A genome 170 unique alleles were identified, 144 unique alleles were identified in the B genome, and 174 unique alleles were identified in the D genome (**Figure 4**, **Table 1** and **Table S4**). In several instances, lines were identified that contained more than one induced SNP in the B or D homoeologs (**Table S4**). The wheat PBF gene consists of two exons separated by an intron of approximately 1 kb. The first exon consists of an untranslated sequence, while the entire protein coding sequence of the gene is present in the second exon [50]. We identified lines containing novel alleles primarily in the N-terminal zinc-finger DNA-binding region, although in the case of the A and D genome homoeologs, we also identified novel alleles in the C-terminal region, which is also thought to play a role in DNA binding and in protein-protein interactions [57]. In addition to identifying premature stop mutations, we also identified missense mutations that were expected to inactivate the WPBF transcription factor by preventing the formation of the zinc finger due to alterations of the cysteines that coordinate the zinc atom necessary for DNA binding. Similar mutations of zinc-coordinating cysteines in DOF genes isolated from pumpkin and maize have been created by site-directed mutagenesis and the corresponding proteins have been shown to lose their DNA binding ability using gel shift and southwestern blotting methods [58] [59] [60]. The totality of the novel alleles identified in these WPBF homoeologs provides a rich resource for further analysis that will help to determine critical residues for their function.

**Figure 4.**
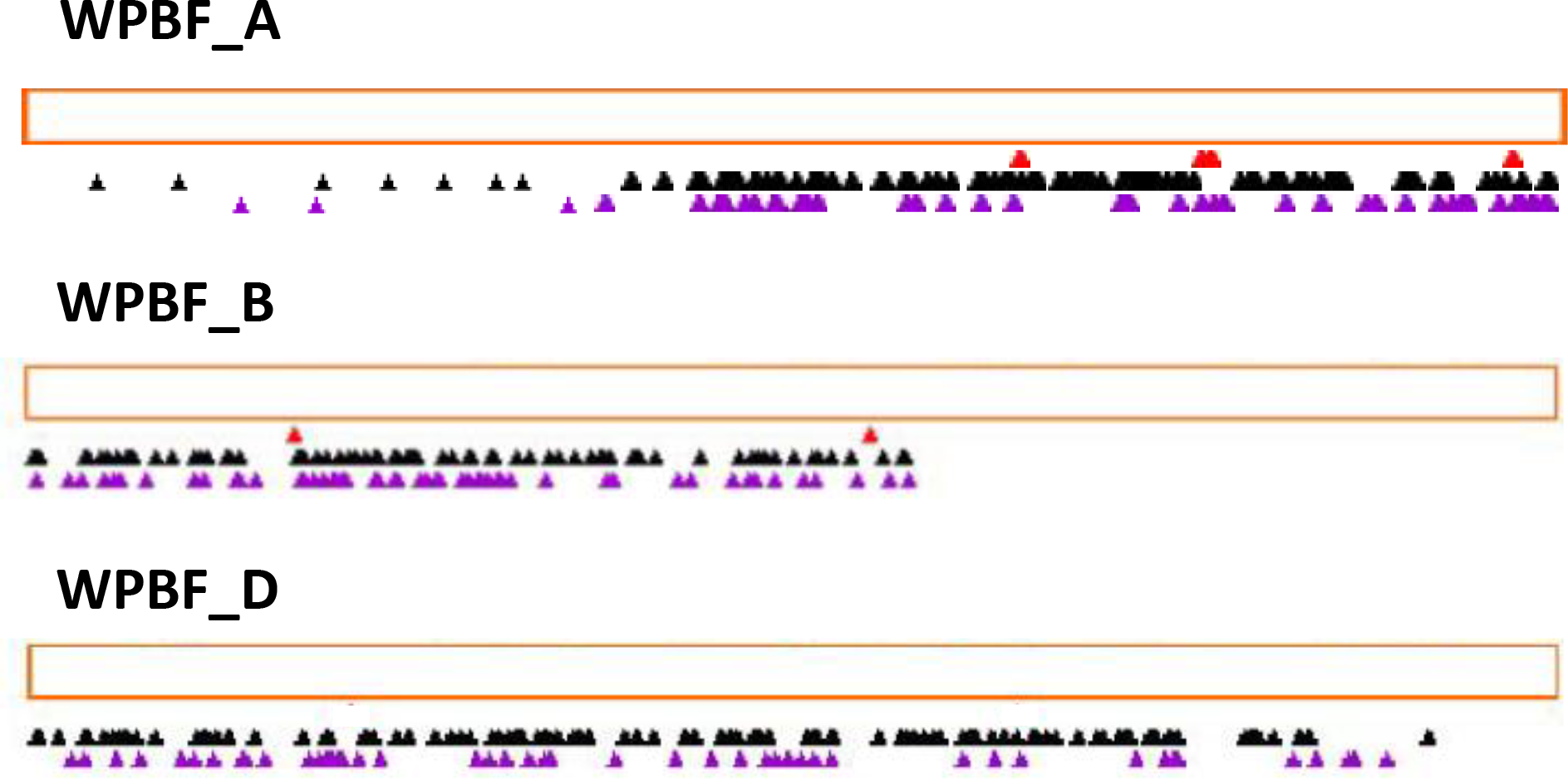
Overview of novel TILLING alleles found in WPBF homoeologs. The red-outlined rectangles represent the A, B and D genome coding regions of the WPBF homoeologs, and the arrowheads show the locations of the TILLING alleles. Red arrowheads represent premature stop alleles, while black arrowheads refer to missense alleles, and purple arrowheads represent silent alleles. Depiction of mutants was made using the program PARSESNP [84].

**Table 1.** Numbers of novel TILLING alleles in A, B, and D genome WPBF homoeologs. Numbers of TILLING alleles of the different classes in wheat are listed. Numbers are total alleles, with numbers of unique, non-redundant alleles in parentheses.

|  | non-coding | silent | missense | severe missense | nonsense | total |
| --- | --- | --- | --- | --- | --- | --- |
| **WPBF_A** | 37 (24) | 65 (42) | 126 (72) | 37 (27) | 8 (5) | 273 (170) |
| **WPBF_B** | 6 (3) | 104 (55) | 82 (43) | 68 (40) | 4 (3) | 264 (144) |
| **WPBF_D** | 18 (13) | 74 (48) | 100 (69) | 60 (44) | 0 | 252 (174) |
| **total** | 61 (40) | 243 (145) | 308 (184) | 165 (111) | 12 (8) | 789 (488) |

### Effects on Protein and Amino Acid Accumulation Due to Inactivating Mutations in WPBF Homoeologs

Because of the hexaploid nature of bread wheat and the expectation that WPBF-inactivating mutations, like the *lys3a* BPBF mutant, would be genetically recessive, we crossed wheat plants containing mutations in the A, B, and D genomes to each other to establish lines containing putative inactive WPBF homoeologs in all three genomes. The first lines we chose to cross to each other included one containing a premature stop mutation in the WPBF gene in the B genome (W70*), and two lines containing tyrosines in place of zinc-coordinating cysteines in the A (C66Y) and D (C63Y) genomes. These were chosen because they were the first TILLING alleles identified in each of the three homoeologs that putatively inactivated the WPBF. Additional severe and, in the case of the A genome, premature stop mutations, were identified later. **Figure 5A** depicts the results of SDS-PAGE analysis of storage proteins extracted from individual seeds of the parental control Express cultivar and from individual F3 seeds, each derived from an independent individual F2 triple null seed. Strikingly, each triple null homozygous seed exhibited an overall decrease of about 50-60% in the accumulation of a variety of gliadins and low molecular weight glutenins as assessed by quantitative analysis of proteins extracted from individual seeds (**Figure 5B**). No decrease in seed storage protein accumulation was observed in F3 seeds derived from triple wild type F2 segregants lacking the mutations (**Figure 6A**). Highlighting the recessive nature of the mutations and the redundancy of the WPBF homoeologs, **Figure 6A** also shows that there is no storage protein reduction in individual F3 seeds from segregants in which one un-mutated WPBF homoeolog is paired with two mutated homoeologs. The data demonstrate that a single wild-type copy on any of the sub-genomes can compensate for missing function on any of the other two sub-genomes.

**Figure 5.**
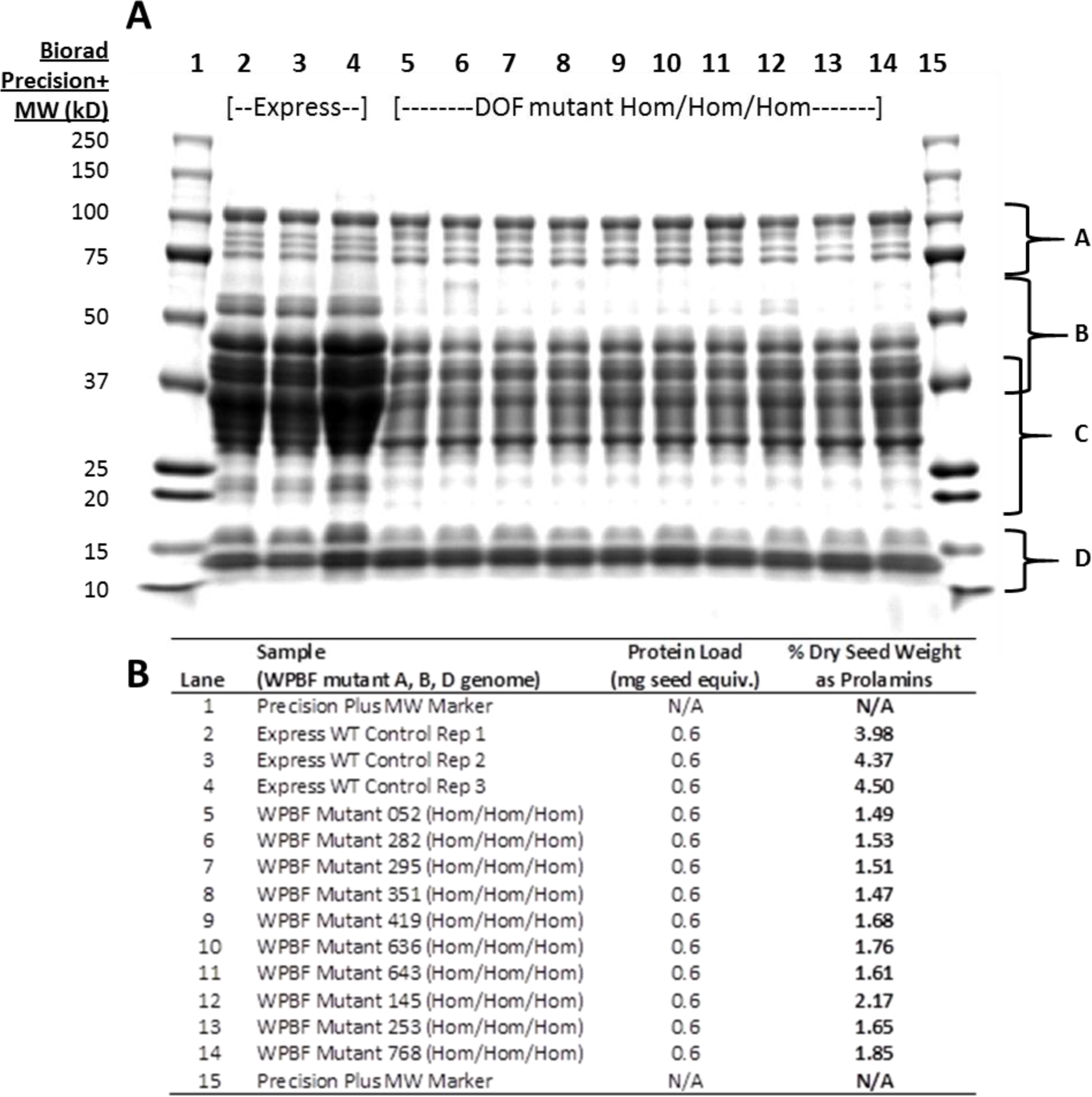
SDS PAGE of wheat prolamins from parental cultivar Express and from WPBF mutants. Panel A depicts a 12% SDS-PAGE gel of seed extracts from parental wheat cultivar Express (lanes 2-4) and lanes 5-14 represent seed extracts from lines in which all three homoeologs of WPBF are inactivated. Each lane shows proteins derived from a single F3 seed and each F3 seed is harvested from a separate F2 plant. Regions of the gel labeled A through D refer to different seed storage protein fractions. A are the HMW glutenins; B are LMW glutenins; C are alpha, alpha-beta, and gamma gliadins; and D are gliadin/avenin-like proteins as well as trypsin and alpha amylase inhibitor proteins. Panel B lists the amount of prolamins in each F3 seed as a percentage of dry seed weight, determined as described in the text.

**Figure 6.**
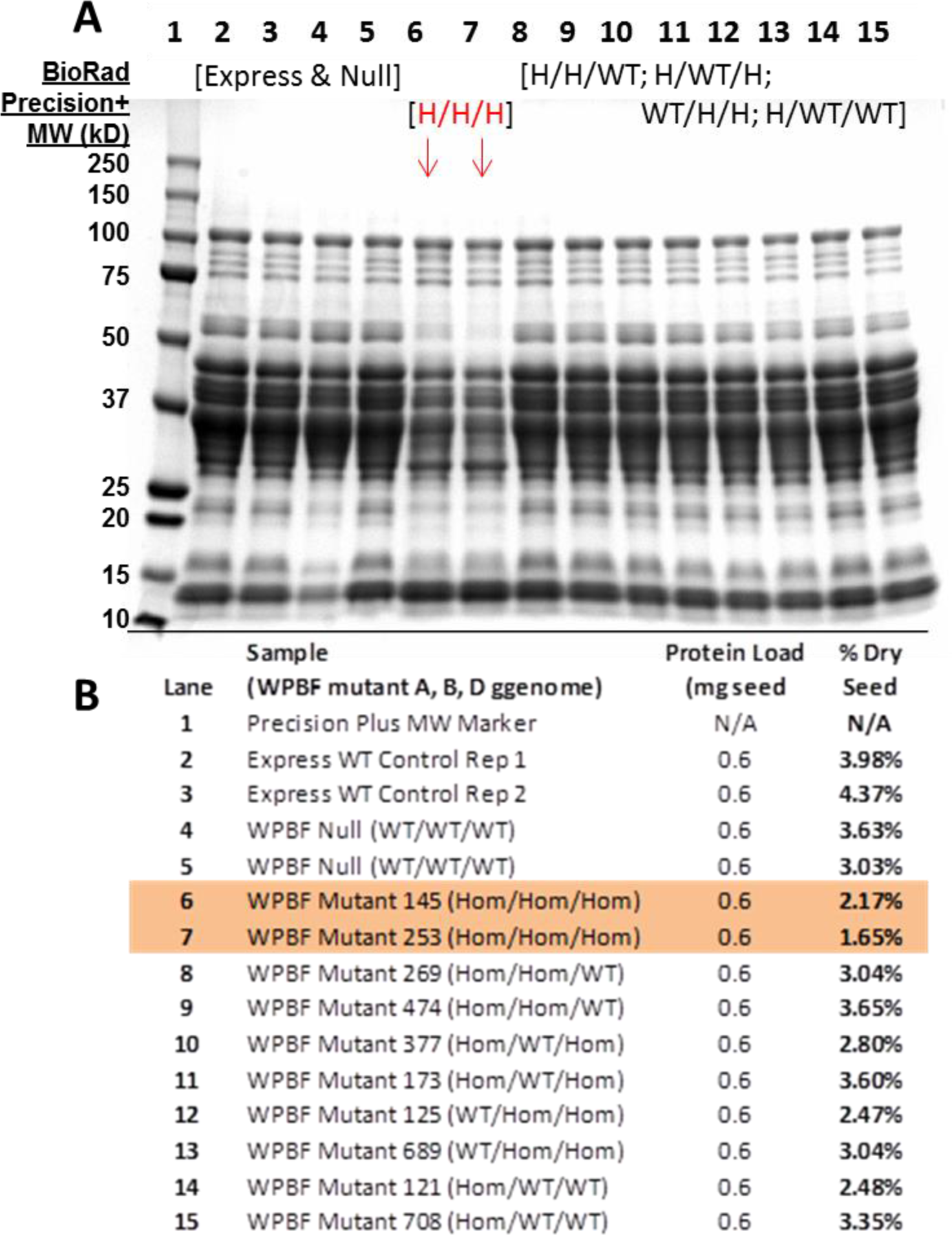
SDS PAGE of wheat prolamins from multiple genotypic classes of WPBF mutants. Panel A depicts a 12% SDS-PAGE gel of seed extracts from parental wheat cultivar Express (lanes 2,3), sibling wild type segregants (lanes 4,5), triple WPBF inactivating alleles (lanes 5,6), and genotypic classes in which one or two homoelogs are not mutant (lanes 8-15). Panel B lists the genotypes in each lane and the amount of prolamins in each seed as a percentage of dry seed weight.

The *lys3a* barley mutant notably fails to accumulate C hordeins and has drastically reduced accumulation of B hordeins. To assess which proteins are absent or reduced in the WPBF triple null mutants, we conducted mass spectrometry (LC-MS) of excised SDS-gel fragments **(Table S5)**. Bands that fall within the region of 30-55 kDa, which make up approximately 70-80% of the total wheat seed storage proteins, are expected to contain α and γ gliadins as well as LMW glutenins, and this was confirmed by the putative identities of the proteins we identified by LC-MS in gel bands excised from this region of the gel **(Table S5)**. Two prominent bands at about 55 kDa, which were drastically reduced in the triple null line were identified as beta-amylase and a LMW glutenin protein [61]. In general, the gliadins and LMW glutenins that are reduced in accumulation in wheat by the triple null WPBF mutations are homologous to the B and C hordeins whose accumulation is reduced by the *lys3a* mutation in barley. These results provide evidence that the A and D genome C>Y mutants are inactivating mutations, as expected, and provide additional evidence that the barley mutation in the BPBF gene underlies the *lys3a* phenotype.

In addition to affecting seed storage protein accumulation, the WPBF triple null line also alters amino acid accumulation in seeds. We extracted total amino acids from seeds of three independent triple null F3 lines, each derived from an independent F2 seed, and compared the content of free amino acids in these seeds to three independent triple wild type segregants. As shown in **Figure S8 and Table S6**, the triple null wheat lines contain considerably reduced amounts of proline, glutamate (deamidated glutamine) and phenylalanine as expected based on their lower levels of gliadins and LMW glutenins, which contain repetitive sequences rich in these amino acids. Decreases in these amino acids were also observed in *lys3a* barley [62]. Among the essential amino acids, most notable is that the triple null lines exhibit 33% and 14% significantly increased amounts of lysine and threonine, respectively, when compared to their wild type siblings. These two amino acids are generally limiting for human nutrition in cereals; efforts have been made to increase lysine in other cereals, and this has led to the development of quality protein maize (QPM), for example. Researchers have concluded that QPM provides health benefits in undernourished populations [63] [64].

## Discussion

We describe the first lowered gluten, high lysine wheat developed with the non-transgenic mutation breeding technique known as TILLING. These lines contain inactivating mutations in the three homoeologs of WPBF, which is a transcription factor that plays a regulatory role in the accumulation of seed storage reserves in cereals. This scientific advance was enabled by our identification of a missense mutation in BPBF as the genetic basis of the low hordein *lys3a* barley mutant. Prior to our wheat TILLING results, no null mutations were known in PBF homologs in any cereal, a fact which had led to the speculation that null mutations might be inviable [65].

Using BSR-seq and genetic fine mapping, we determined that the *lys3a* missense mutation is responsible for the pleiotropic defects found in *lys3a* barley. In addition to a drastic reduction in the accumulation of B and C hordeins [66], this mutant exhibits multiple other effects including an increase in the accumulation of free and protein-bound lysine [18], an increase in the embryo size [67], a reduction in starch accumulation as well as shrunken seeds and a decrease in yield [68]. Concerted breeding efforts, whose objective was to increase yield while maintaining the high lysine phenotype, without regard to the hordein phenotype, however, minimized the negative effects of the mutation. Due to the linked nature of the high lysine and low hordein phenotypes, these efforts also maintained the low hordein phenotype [68]. Given the major effects of the *lys3a* mutation, we hypothesize that the missense mutation we identified in the DNA binding domain of the BPBF transcription factor likely inactivates it. Experiments to investigate the DNA-binding properties of this BPBF mutant using electrophoretic mobility shift assays were unsuccessful (data not shown); however, transgenic expression of wild type BPBF partially complemented the *lys3a* mutation, which supports our identification of this gene as the basis of the *lys3a* mutation.

The results of the BSR-seq analysis conclusively showed that the *lys3a* mutation maps to the pericentromeric region of chromosome 5H (previously known as chromosome 7). This is consistent with earlier mapping experiments [24] [16]. Nevertheless, in a recent paper describing the systematic backcrossing and mapping of a large collection of barley mutants [26], the *lys3a* mutant was mapped to chromosome 1H (line ID: BW496) [27]. We obtained the described putatively *lys3a*-containing line BW496 [26] from the USDA Small Grains Germplasm Collection and extracted hordeins from endosperms of multiple seeds. SDS-PAGE of hordeins appeared to be normal with none of the obvious, characteristic hordein alterations found in the *lys3a* mutant (data not shown). We conclude that this mutation was not present in the seeds of line BW496 that we obtained.

Due to studies by Sørensen [69] showing that B-hordein promoters remained methylated in the endosperm of *lys3a* mutants while B-hordein promoters were unmethylated in the wild type endosperm, it was proposed that the *lys3a* mutation might be in a 5-methyl-cytosine glycosylase (Demeter) enzyme that removes methyl groups from methylated DNA [70]. However, the main barley Demeter homolog is found at approximately 400Mb on the long arm of chromosome 5H, a considerable physical distance from the *lys3a* mutation. Nevertheless, it is apparent that DNA methylation also plays a role in affecting seed storage protein accumulation in cereals [71] [72] [70]. The possible connection between PBF genes and DNA methylation in cereal endosperms merits further investigation.

Although the degree of prolamin reduction in our current wheat PBF TILLING lines (50-60%) is not sufficient to make them appropriate for celiac patients, additional TILLING approaches are underway that we anticipate will reduce gluten even further. In addition, the wheat PBF TILLING lines may be combined with other approaches, such as gluten-degrading enzyme supplements [73] [74], to benefit individuals with gluten sensitivities. In our previously published work, we provided evidence for this synergistic approach using a non-human primate model of celiac disease [75]. We showed that the *lys3a* barley mutant reduces prolamin-related symptoms when it replaces conventional barley in the feed given to gluten sensitive rhesus macaques [76]. Furthermore, symptoms due to prolamin consumption in these animals were not observed when this low hordein barley feed was supplemented with a prolamin-degrading enzyme [77]. Future research using reduced gluten wheat will be necessary to expand on these findings.

Because the accumulation of the HMW glutenins appeared not to be substantially affected in our WPBF-mutant lines, this reduced gluten wheat may retain some of the important viscoelastic functional properties of normal wheat. We are currently increasing seed of these lines in order to address this question. Because wheat is able to tolerate a high mutation load [80], each TILLING line contains a large number of background mutations that must be removed by genetic crosses. Like the barley *lys3a* mutant, the triple homozygous mutant wheat lines described exhibit some negative pleiotropic effects such as smaller seed size **(Figure S9)** and somewhat reduced total protein and starch levels **(Table S7)**. Future breeding will be required to determine if these effects can be overcome. Due to incompletely understood mechanisms of seed proteome rebalancing [81], the reduced gluten wheat lines we developed also contain higher levels of lysine and threonine, making them more nutritious for all consumers of wheat and products made with wheat. Using the molecular markers developed in the present study for marker assisted breeding, the potential agronomic and functional utility of these wheat lines can be determined.

## Acknowledgements

For a gift of seeds of barley cultivar “Lysiba” and for friendly collaboration during the early stages of this research the authors thank Dr. Diter von Wettstein (deceased). The authors thank Dr. Harold Bockelman at the USDA National Small Grains Collection in Aberdeen, ID for seeds of Bomi, *lys3a* barley (Risø 1508), BW496, and the barley iCore. For excellent technical assistance, the authors are grateful to Ennis Sandle and Yingzhi Lu. For critical reading of the manuscript the authors are grateful to Dr. Ann Slade and to Dr. Margaret Miller. This work was funded by the National Institute of Diabetes and Digestive and Kidney Diseases of the National Institutes of Health under Award Numbers R42DK097976 and R42DK072721. The authors alone are responsible for the content of this article and this work does not necessarily represent the views of the National Institutes of Health.

## Author Contributions

C.P.M. wrote the manuscript and performed and coordinated research and was the PI on the NIH grants. W.J.A. performed half-seed analysis, SDS PAGE, construct development, genotyping and molecular research. A.H., D.L. and J.M performed TILLING assays and genotyping. T.A.G.L. performed molecular research. P.S.S. and his staff at Data2Bio LLC conducted the BSR-seq. W.S. conducted mass spectroscopy and protein and amino acid analysis. J.v.B. coordinated protein and transformation research. L.W. was responsible for barley transformation. C.M. coordinated research and conducted wheat crosses and analysis. All authors read and approved the manuscript.

